# Evolution and Functional Information

**DOI:** 10.1101/114132

**Authors:** Matthew K. Matlock, S. Joshua Swamidass

## Abstract

“Functional Information”—estimated from the mutual information of protein sequence alignments—has been proposed as a reliable way of estimating the number of proteins with a specified function and the consequent difficulty of evolving a new function. The fantastic rarity of functional proteins computed by this approach emboldens some to argue that evolution is impossible. Random searches, it seems, would have no hope of finding new functions. Here, we use simulations to demonstrate that sequence alignments are a poor estimate functional information. The mutual information of sequence alignments fantastically underestimates of the true number of functional proteins, because it also is strongly influenced by a family’s history, mutational bias, and selection. Regardless, even if functional information could be reliably calculated, it tells us nothing about the difficulty of evolving new functions, because it does not estimate the distance between a new function and existing functions. The pervasive observation of multifunctional proteins suggests that functions are actually ver close to one another and abundant. Multifunctional proteins would be impossible if the FI argument against evolution were true.

## 1 Introduction

Evolution is among the most successful theories in all of science. It rationalizes a staggering array of otherwise disparate biological observations. In our genomic age, the evidence for evolution is more clear than ever before. Many independent patterns in our genomes point to the common ancestry of life. Despite the broad scientific consensus around common ancestry, evolution remains controversial to the general public, and is a focal point of ongoing political challenges [17, 18].

In this societal context, it is important to correct faulty arguments against evolution [7, 25]. This is particularly important when components of these arguments appear in peer reviewed journals. To this end, this study aims to explain the modeling errors on which the “Functional Information” (FI) argument against evolution depends. Components of this argument are published in several peer reviewed articles [8, 13], and are elaborated in public debates, rebuttals and counter rebuttals (see SI). Science, of course, is not litigated through popular debates, but in careful studies. So it is important the faulty scientific claims in peer-reviewed publications are corrected, which is what we aim to do here.

In the following sections, we first explain the FI argument. Next, we demonstrate with evolutionary simulations that the key scientific claims on which this argument rests are false. To the point, sequence alignments wildly overestimate FI and wildly underestimate the number of functional sequences. Finally, we explain why FI is not a valid way of estimating the difficulty of evolving new functions, even if FI could be reliably estimated from sequence alignments.

### 1.1 The Functional Information Argument

Functional information (FI) itself is a legitimate concept introduced in the literature to quantify and study the emergency of complexity in evolutionary systems [13]. The FI argument against evolution attempts to develop estimates of functional information and to use these estimates to show that complex biological systems are highly improbable to be the products of evolution.

The FI argument is formulated in multiple ways. In one formulation, FI is estimated by computing the mutual information of a sequence alignment (MISA)—which is sometimes termed functional sequence complexity (FSC)—of a family of proteins with a specified function. Next, it is hypothesized (without verification) that intelligent minds are uniquely capable of producing significant amounts of FI, more than 150 bits of MISA. Then, it is asserted that alignments of proteins with common function demonstrate that biological function is characterized by very high MISA. Therefore, we must conclude that life must have been designed, and could not have evolved from random processes.

In a parallel formulation, MISA is used to estimate the number of proteins of a given length with a specified function, by assuming that MISA is a good estimate of FI. From here, it is asserted that the chances of finding a protein with a specified function is astronomically small if the measured FI is greater than 150 bits. Therefore, the argument concludes that it is impossible for proteins with the specified function to have evolved by random chance.

Both variations of the argument rely on a similar foundation: the assertions that (1) MISA is an accurate estimate of FI and that (2) FI is a meaningful measure of evolvability. In the literature and the public debates, MISA and FI are often conflated and treated as though they are the same quantity (SI Data). Keeping these concepts separate, and treating FSC as a synonym for MISA, clarifies the scientific questions at hand; is MISA (i.e. FSC) a good estimate of FI? And is FI, even if it were knowable, a good measure of evolvability?

### 1.2 Functional information and the probability of function

Following from information theory, we define the “functional information” of a specific biological function as

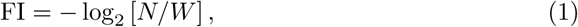

where N is the number of protein sequences with the specified function and W is the total number of possible protein sequences under consideration [13]. From this formula, the probability of guessing a functional sequence decreases exponentially as the FI increases (2^-FI^). This interpretation only holds if each sequence is guessed with equal probability.

### 1.3 Functional information from sequence alignments

The central scientific claim: the MISA of proteins with a specified function is a good estimate of the function’s FI. This claim is so confidently asserted that MISA and FI are sometimes conflated and treated as though they are the same quantity. This claim appears in the peer-reviewed literature, authored together by a group of Intelligent Design theorists [8]. They coin a new term FSC, but use the formula for mutual information to compute it,

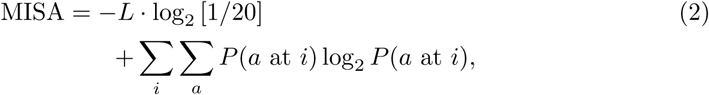

where *L* is the length of the sequence alignment, *i* ranges over each position in the alignment, *a* ranges over the twenty amino acids, and *P(a* at *i*) is the probability of observing amino acid *a* at position *i* [8]. This formula applies to both position specific scoring matrices (PSSMs) and Hidden Markov Models (HMMs), and it is asserted that Pfam—the widely used repository of protein family HMMs [9]—can be used to estimate the FI of important biological functions.

Care is given to remove redundant sequences and ensure that enough sequences are included for the MISA to converge to a specific value. Measured this way, the MISA of important protein families ranges from about 1 to 4 bits per amino acid. MISA is computed to be over 150 bits per family for many protein families, and up to 1,000 bits for some families.

### 1.4 Functional information and evolvability

The final step of the FI argument combines equations 1 and 2 to calculate the probability of randomly guessing a protein sequence with a specified function. Previous studies estimate that evolution has explored at most 2^145^ different protein sequences [7]. Therefore, the argument concludes that any protein function we observe with substantially more than 150 bits of FI (measured by MISA) is very unlikely to have evolved. In particular, the argument claims that protein families with greater than 1000 bits MISA are astronomically unlikely to be the products of evolution.

## 2 Results and Discussion

The FI argument depends on several fallacies, but we focus on the component of the argument that appears in the literature. Specifically, straightforward simulations demonstrate that MISA wildly overestimates FI. Furthermore, we argue that FI (even if accurately estimated) cannot be used to quantify the difficulty of evolving a specific function. Proceeding forward, it is important to emphasize a sharp distinction between the FI estimated by MISA (Equation 2) and the true FI computed from the total number of known and unknown sequences with the specified function (Equation 1). As we will see, there is a fantastically large gap between MISA and the true FI.

### 2.1 Uniform evolution of non-functional proteins

This first and most important simulation demonstrates that mindless evolutionary processes can produce sequences with very high MISA. This simulation samples several extant sequences a fixed number of mutational events away from a single ancestral sequence. There are no functional constraints, so the FI by definition equals zero (Equation 1). If MISA is a good estimate of FI, it should converge to zero bits too.

We find that MISA does not converge to zero, even as more extant sequences are sampled (Figs. 1 and 2). Entirely mindless processes produce sequences with high, mutation-rate- and length-dependent MISA. The simulation converges to 4.3, 2.6, 1.8, 1.2, and 0.8 bits per amino acid for mutation rates of, respectively, 0.0, 0.25, 0.50, 0.75, and 1.0 mutation events per amino acid. MISA does go to zero if the mutation rate is increased to several events per amino acid, as the memory of the ancestral sequence is wiped. So, in addition to the effect of functional constraints, MISA encodes the mutational age of the protein family.

**Figure 1.**
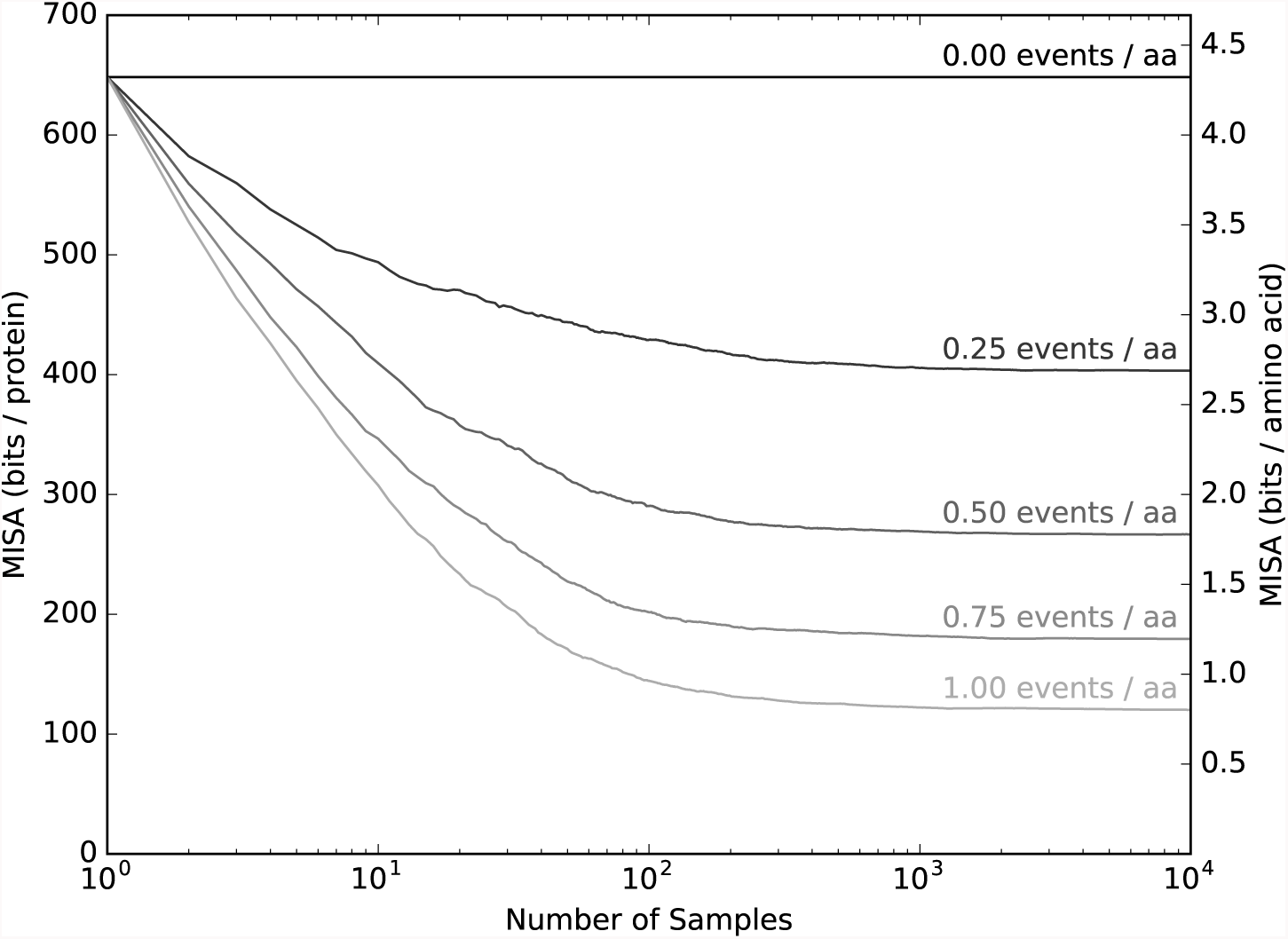
MISA is arbitrarily high even for sequences with an FI of zero. Here, extant sequences 150 amino acids long, and a fixed number of mutations away from a single ancestral sequence are sampled, added to the sequence alignment, and the MISA is reported as a function of the number of samples

**Figure 2.**
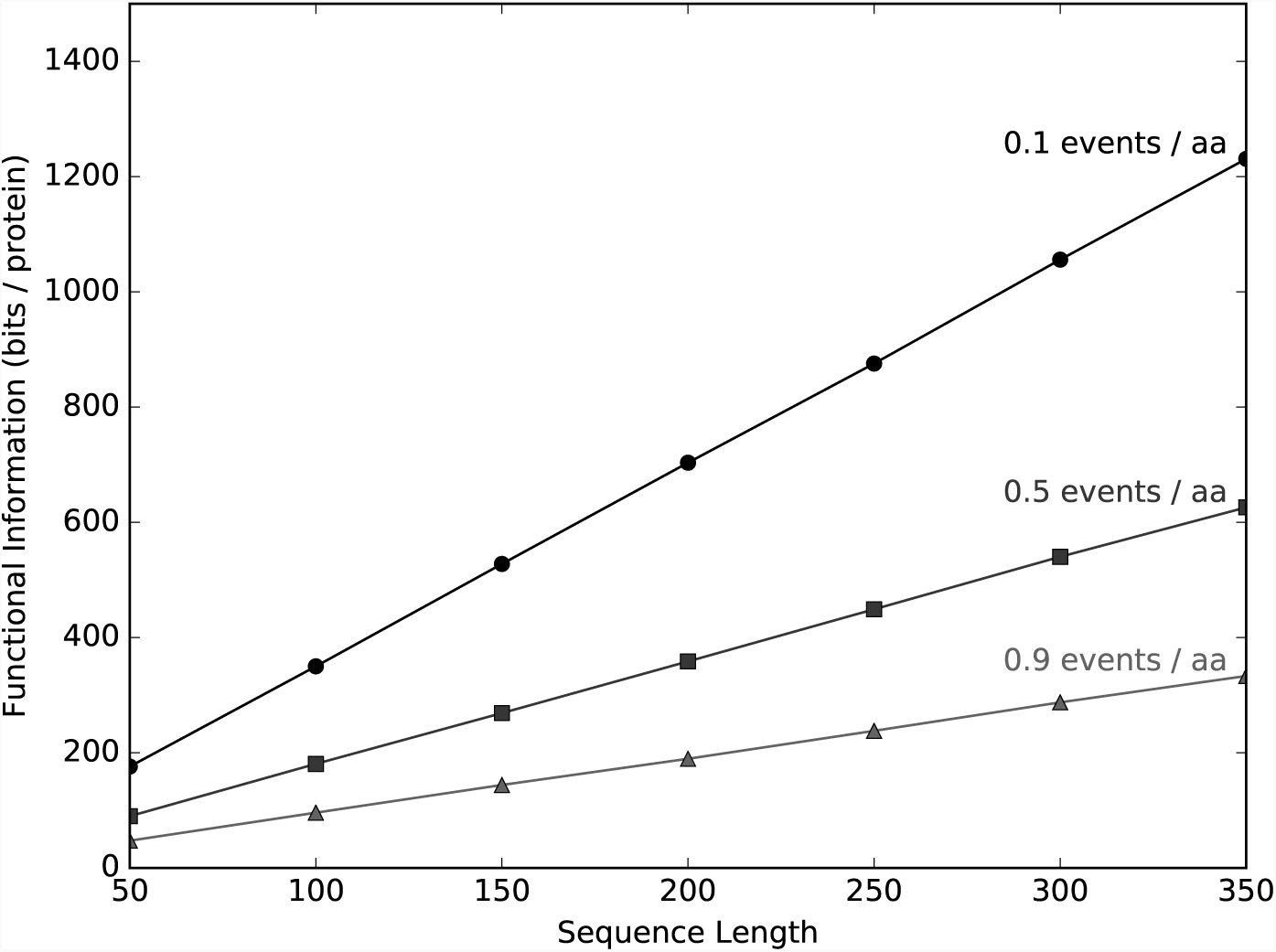
MISA increases scales linearly with sequence length, even when true FI is zero. Here, extant sequences 150 amino acids long, and a fixed number of mutations away from a single ancestral sequence are sampled, added to the sequence alignment, and the MISA is reported as a function of the number of samples. The MISA of this process scales linearly with sequence length, even though the true FI is zero

The mutation rates in the simulation correspond to very long time periods. One mutation per amino acid corresponds to more than 30 million years of mammalian evolution (at approximately ten mutations per base per billion years). Moreover, this effect would amplified by sampling extant sequences from a phylogenetic tree, where some sequences more recently diverge from one another.

Therefore, the MISA of protein families is strongly influenced by the common evolutionary origin of the extant sequences. Evolution samples sequences by randomly walking around the ancestral sequence, not by randomly sampling sequences with equal probability from the space of all possible proteins. This biased sampling pushes MISA upward. The effect of evolutionary history on MISA is substantial, protein length dependent, and independent of functional constraints.

As obvious as this result may seem, its importance in this discussion cannot be overstated. This simulation directly falsifies the hypothesis that “minds are uniquely capable” of generating sequences with high FI as measured by MISA. From this experiment, we must conclude either that (1) a mindless process driven entirely by randomness can reliably generate arbitrarily high FI or that (2) MISA is an extremely bad estimate of FI. Either conclusion falsifies the FI argument entirely.

Going forward, any sensible effort to measure FI from a sequence alignment needs to subtract the effect of shared evolutionary history. Put another way, the information in a sequence alignment (MISA) quantitates both functional and historical information. The FI encodes a sequence’s function, but the historical information records the sequence’s history, its ancestry. Any effort to compute FI needs to subtract the historical information from MISA. One solution to this problem might use DNA sequences adjacent to protein coding regions as a control. The mutual information of these sequences, if not under strong functional constraint, might be used to compute a length dependent correction factor. Full consideration of this possibility is beyond the scope of this study. However, if done correctly, an appropriate correction would substantially reduce estimates of FI in many cases. Regardless, MISA itself wildly overestimates FI.

### 2.2 Codon evolution of non-functional proteins

In the first simulation, as mutation rate increases to several events per amino acid, the substitution rate approaches 100% and the MISA of the extant sequences approaches zero bits. This is only true because the simulation uses a mutational model that uniformly substitutes amino acids. Biological systems, however, substitute amino acids non-uniformly, and this introduces an additional bias.

A more accurate simulation models the evolution of codons with restrictions imposed by the genetic code. Here, an ancestral sequence is constructed by uniformly sampling codons. Mutations change a codon into another codon with exactly one nucleotide difference. As would be expected, the MISA of non-functional sequences generated by codon evolution is even higher than that generated with a uniform model. The simulation converges to 4.3, 3.2, 2.5, 2.1, and 1.6 bits per amino acid for mutation rates of, respectively, 0.0, 0.25, 0.50, 0.75, and 1.0 mutations per amino acid (Fig. 3). Once again, MISA scales inversely with mutation rate. In this model, the equilibrium amino acid composition is not uniform but is proportional to the number of codons encoding each amino acid. Consequently, even after a large number of mutations erase all memory of the ancestral sequence, an additional 0.183 bits per amino acid are added to MISA, and it never reaches zero (Fig. S1).

**Figure 3.**
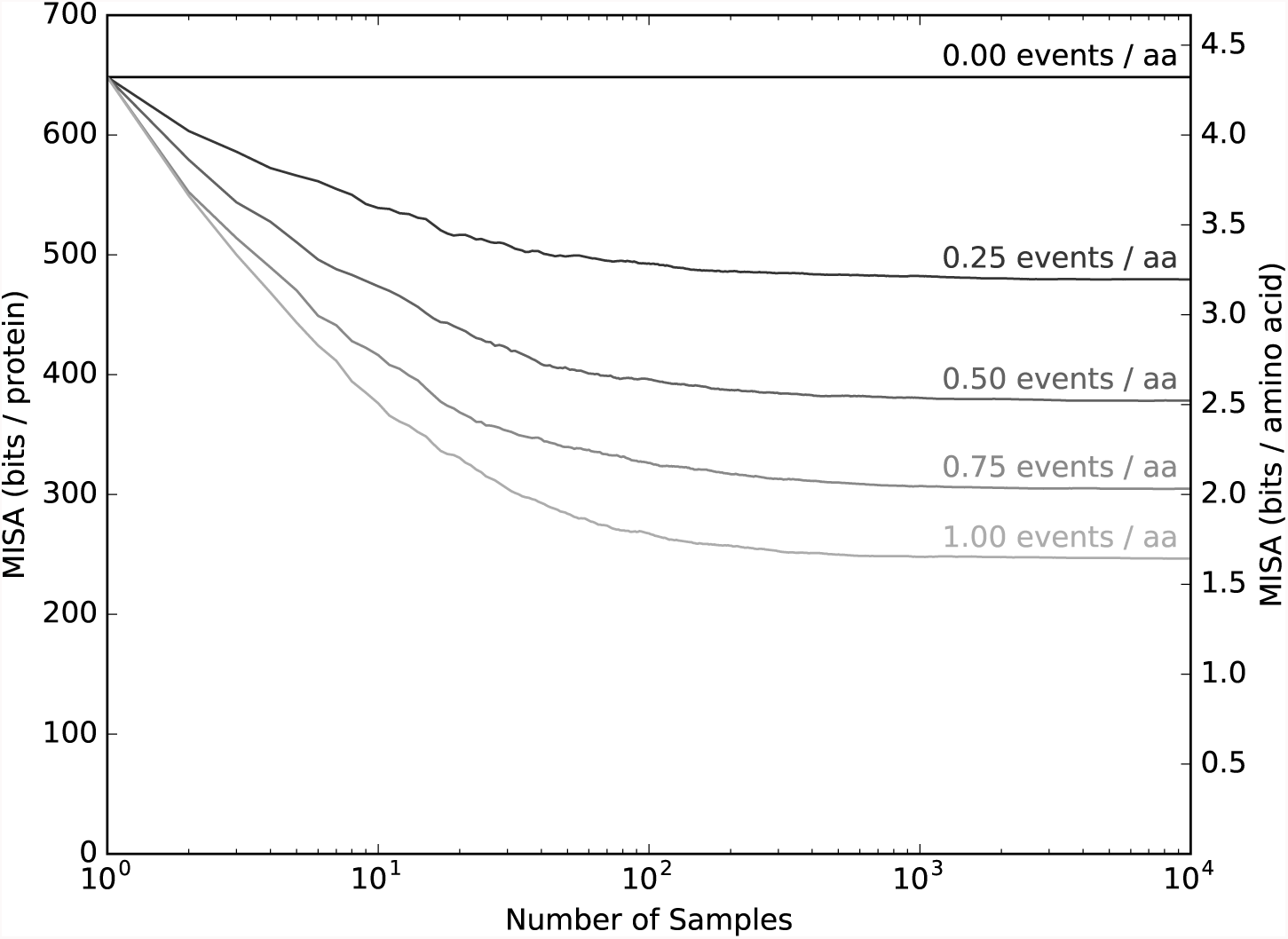
Codon evolution of non functional sequences. The simulation models the evolution of a codon sequence, by limiting mutations to substitution of codons with a single base-pair difference. Extant sequences a fixed number of mutations away from a single ancestral sequence are sampled, added to the sequence alignment, and the MISA is reported as a function of the number of samples. As the number of samples increase, MISA does not converge to zero if the mutation rate is not high enough, even though the true FI is zero. As mutation rate increases, asymptotic MISA does move downwards, but more slowly than uniform evolution and it does not ever reach zero FI even once all amino acids have been substituted

So, improved modeling of mutational mechanisms increases the measured MISA. MISA is only a good estimate of FI if known sequences are uniformly sampled from the space of functional sequences. However, the more realistically evolution is simulated, the less uniformly sequences are sampled, and this deviation from uniformity adds to MISA.

### 2.3 More realistic mutational models

It is beyond the scope of this study to implement a comprehensive model of evolutionary change. Rather, we emphasize that MISA estimates of FI are inaccurate, and simulations are made more realistic, MISA diverges farther from FI. Each improvement in the evolutionary model introduces additional non-uniform biases into the sampling of sequences, which add an additional upward bias to MISA estimates of FI.

More realistic simulations would model several additional mutational mechanisms. For example, genome GC-content also exerts a strong effect on amino acid composition and substitution probabilities [4, 16, 21, 23]. Likewise, differences in transversions and transition probabilities are also important, are caused by biochemical mechanisms, and appear to be independent of amino acid conservation [22]. A comprehensive model would also include insertions, deletions, stop codons, frameshift mutations, exon shuffling, exon duplication and deletion, alternate splicing, recombination, fusions, transposons, horizontal gene transfer, and more. Of course, some of these mechanism would require more complicated alignment algorithms than PSSMs to correctly compute MISA. For example, sequences from a mutational model that includes insertions and deletions would be correctly aligned by a HMM, but not a PSSM. Unfortunately, there are no standard approaches to aligning sequences generated by many important mutational mechanisms. Each mechanism introduces a new source of “information” into protein sequence families. In the same way codon evolution introduces a persistent deviation from uniformity that pushes MISA upwards, so does each one of these mechanisms.

One poorly motivated strategy to cope with the complexity of mutational mechanisms is to settle for alignment methods incapable of correctly aligning sequences. Attempts to justify this approach might incorrectly assert that alignment deficiencies can only bias MISA estimates downward, producing more conservative estimates of FI. However, this assertion is predicated on correct identification and alignment of sequences with the same function. Methods incapable of aligning sequences under a realistic mutational model will introduce upward biases to MISA in at least three ways. First, excluding functional sequences from the alignment reduces the sampling of the alignment, which in many substitution regimes increases MISA estimates (Fig. 1). Second, alignment deficiencies makes it much harder to identify all the families with shared function. Third, failure to align to correctly functional sequences multiplies the number of sequence families with the function, and the more functional families there are, the more divergent MISA is from FI (as we will see).

Unfortunately, no method is guaranteed to correctly align sequences generated by all these mutational mechanisms. For example, there is no standard method of accounting for the non-linear alignments produced by fusions and exon shuffling. However, speculatively speaking, it is possible that estimates of mutual information computed by DNA compression algorithms might be more accurate than HMMs in some cases; for example, fusions and exon shuffling are not modeled by Pfam HMM alignments but are correctly modeled in genome compression algorithms [5], and compression size estimate an upper bound on information content. Putting the technical challenges aside, the particulars of many mutational mechanisms drive evolution farther from uniform sampling. Therefore, each mechanism, when correctly modeled, can increase estimates of MISA. In fact, unaccounted for patterns in genomic data is exactly how some mutational mechanisms are identified in the first place [3, 11, 12]. Each mutational mechanism has the potential to introduce an independent upward bias to MISA estimates of FI. Each of these biases needs to be identified and accounted for if MISA can ever effectively measure FI.

### 2.4 Multiple families with shared function

The MISA approach to estimating FI assumes that only a single family of proteins can perform a specified function. Even if all the biases presented in this study could be corrected, this assumption fundamentally limits any sequence alignment based approach to estimating the number of proteins with a specified function that are *also* within the same family. This assumption is not warranted. Sequences that cannot be aligned can still adopt the same tertiary structure [15] and can share the same function [19]. A more accurate method would, instead, compute the FI of each individual functional family, and then aggregate them to compute the true FI. It follows algebraically from Equation 1 that the MISA of each family should be aggregated by summing in negative exponential space,

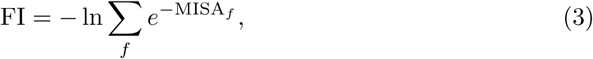

where *f* indexes functional families, and MISAf is the FI of each family. In the toy case, where all families have nearly equal FI, the aggregation reduces to,

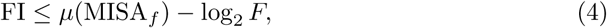

where μ (MISA_*f*_) is the mean MISA across all functional families, and F is the total number of families. This is an upper-bound, because sums in negative exponential space are dominated by smaller terms. In contrast, the mean equally weight all terms, thereby biasing the FI approximation upwards, so this formula is an upper-bound, not an unbiased estimate. Alternatively, this aggregation has another algebraically derived upper-bound,

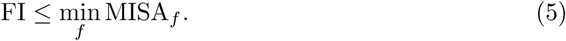

So the MISA of one family is at best a very loose, low confidence, upper bound on FI. Put another way, these results demonstrate the FI of a specified function is always less than the MISA of the simplest protein family with the function. Likewise, it is always less than the average MISA of families with the function. Most important, aggregate FI can be much smaller than the MISA of any individual family with the function.

From this analysis it might seem possible to estimate FI by aggregating MISA across the multiple families that we know have shared function. However, this approach assumes that all families with a specified function have been identified and sequences are available for each family in large numbers. However, evolution does not uniformly sample sequence space, so there is no reason to think we have catalogued all possible families of proteins, or that all of them exist in nature. As has been laid out in the FI argument itself, there is a upper-bound of about 2^145^ protein sequences explored by evolution. But there are 20^150^—or approximately 2^648^—proteins that are 150 amino acids long. To be sure, it is possible that evolution could have explored all possible sequences of shorter proteins, especially if early life had a more limited amino acid alphabet [7]. Nonetheless, it is clear, for the current alphabet of 20 amino acids and proteins with 100 or more amino acids, only an infinitesimally small proportion of protein sequences have been explored. So only a very small number of protein families have been sampled. Simply aggregating the MISA of known families to estimate FI does not manage or correct for this under-sampling bias, and there is no foreseeable theoretical way of overcoming this challenge.

Moreover, we do not even observe (or know the function of) the vast majority of the sequences that evolution has explored. Setting aside the ancient sequences of proteins that have been irrecoverably lost, the recent proliferation of metagenomic studies demonstrate how little we know about extant proteins. In many metagenomic experiments, more than 90% of sequences fail to align to known protein families [6]; the unknown sequences are frequently different from experiment to experiment too. We only know the functional role of only a tiny fraction of sequences. Some of these sequences, undoubtably, represent novel functions, but how many represent novel families with functions already known? Knowing the MISA of one family, even if it could correctly estimate the size of the family, tells us nothing about the aggregate size of all functional families, nor does it tell us how many there are. MISA does not estimate the aggregate MSIA of all functional families (Equation 3), nor does it estimate the minimum MISA across the families (Equation 5). Therefore, MISA tells us very little about FI and, consequently, the number of proteins that have a specified function.

### 2.5 Selection for high function

Selection also introduces a strong upward bias to MISA estimates. In many cases, high efficiency proteins are selected over low efficiency proteins with the same function. Most skeptics of evolution term this “microevolution,” and readily concede that the iterative refinement of proteins by natural selection is a real and observable process. This widely understood, verified, and accepted mechanism increases MISA.

The essential point is that selective pressures, in many cases, will limit extant sequences to those that are high functioning, excluding those that are low function. So sequence alignment of extant sequences could only hope to estimate the FI (or the number) of high functioning proteins, instead of the much larger space of all protein sequences with the specified function, including the less efficient members of the family. This effect is likely to be strongest in ancient proteins that have been well optimized over long periods of time, and at the same time have outsized impact on fitness. For example proteins functioning in DNA replication, RNA transcription, and protein translation are all ancient and strongly impact fitness. Selection likely confines them to a narrow space of high functioning sequences, even though the space of less efficiently functioning sequences could be much larger.

To demonstrate this effect, we simulated evolution of functional sequences under selection using the codon model of evolution (Fig. 5). In this simulation, we see that MISA is much closer to the FI of high functioning sequences, and tells us nothing about the total number of sequences of any functional efficiency. These results are robust to a large number of variations in the parameters, and any reasonable fitness function that places a selective advantage to high over low function (Fig. S3 and S4). The smoothness of the fitness landscape makes no difference in this simulation, as long the high function space is eventually reached. Critically, MISA is influenced very weakly by the size of the low function space. On the other hand, it is almost entirely determined by the size of high function space.

Although a simplified model of evolution, it recapitulates several key behaiviors. Initially, but only for a short time, positive selection dominates. However, when the high functioning space is reached, the simulation is dominated by neutral mutations and purifying selection to keep the population in high functioning space. Moreover, the simulation exposes an unfounded assumption in MISA estimates of FI, that selection operates at the boundary of functional and non-functional sequences. For extant proteins under purifying selection, this is not the case. Rather, selection here operates at the boundary of high and low function sequences. It not clear how MISA could be corrected to adjust for this effect and, as we have seen, this effect is substantial and persists indefinitely.

### 2.6 Functional information and evolvability

Until now, we have focused on the problems with estimating FI from MISA. Our emphasis here is to correct fallacious claims in the literature that conflate FI and MISA, as if the two quantities are nearly equivalent. In contrast with this claim, we identify several very strong upward biases to MISA that are independent of FI: historical information, mutational bias, and selection. MISA, therefore, is a very poor estimate of FI.

Still, it is worth considering if FI, if it were possible to estimate, would be a good estimate of evolvability. Recall, indispensable to the FI argument is the assertion that FI is a good measure of how difficult it is to evolve a specific function. The argument depends on using FI to measure the difficulty of randomly guessing a functional sequences.

This rationale, however, is flawed because evolution does not sample sequences with uniform likelihood. If we accept the definition of FI presented in Equation 1, FI just tells us the proportion of sequences that are functional. If evolution just sampled uniformly from the space of all proteins, it is possible that FI would be a good estimate of evolvability. However, evolution proceeds by sampling around existing protein and DNA sequences, which are substantially enriched for function. Evolvability is better estimated by measuring the distance between existing sequences and the closest family of functional proteins. FI, however, tells us nothing about how close the first functional protein of a family was to preexisting sequences without that function (Fig. 4).

**Figure 4.**
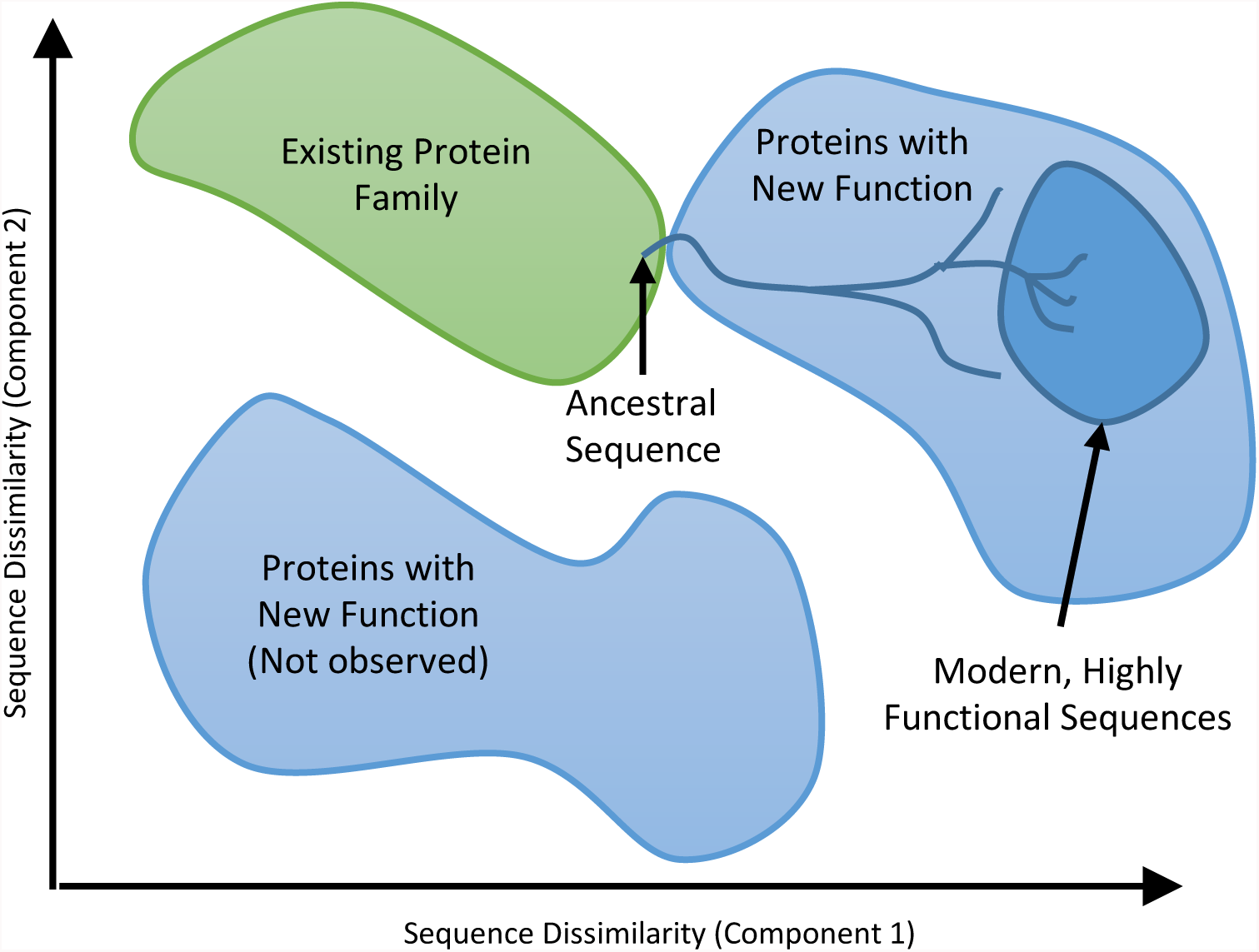
Evolutionary processes cause high MISA in extant sequences, independent of functional constraints. The figure depicts a hypothetical sequence space with regions corresponding to two functions marked. The lines depict the evolutionary trajectory as a new sequence is evolved. Positive selection, in this case, concentrates the extant sequences in the region of highly functional sequences. Estimates of functional information from the sequence alignment of extant sequence are limited to estimating the size of the region occupied by the extant sequences (the dark blue region). Due to incomplete and non-uniform sampling, the full extent of possible functional sequences within the protein family is not correctly measured, and other protein families with the same function are also not counted. The effect of these limitations is that sequences alignments dramatically overestimate functional information and underestimate the number of functional sequences. Regardless, evolvability might be better measured, for example, by the distance between the new function and pre-existing sequences

**Figure 5.**
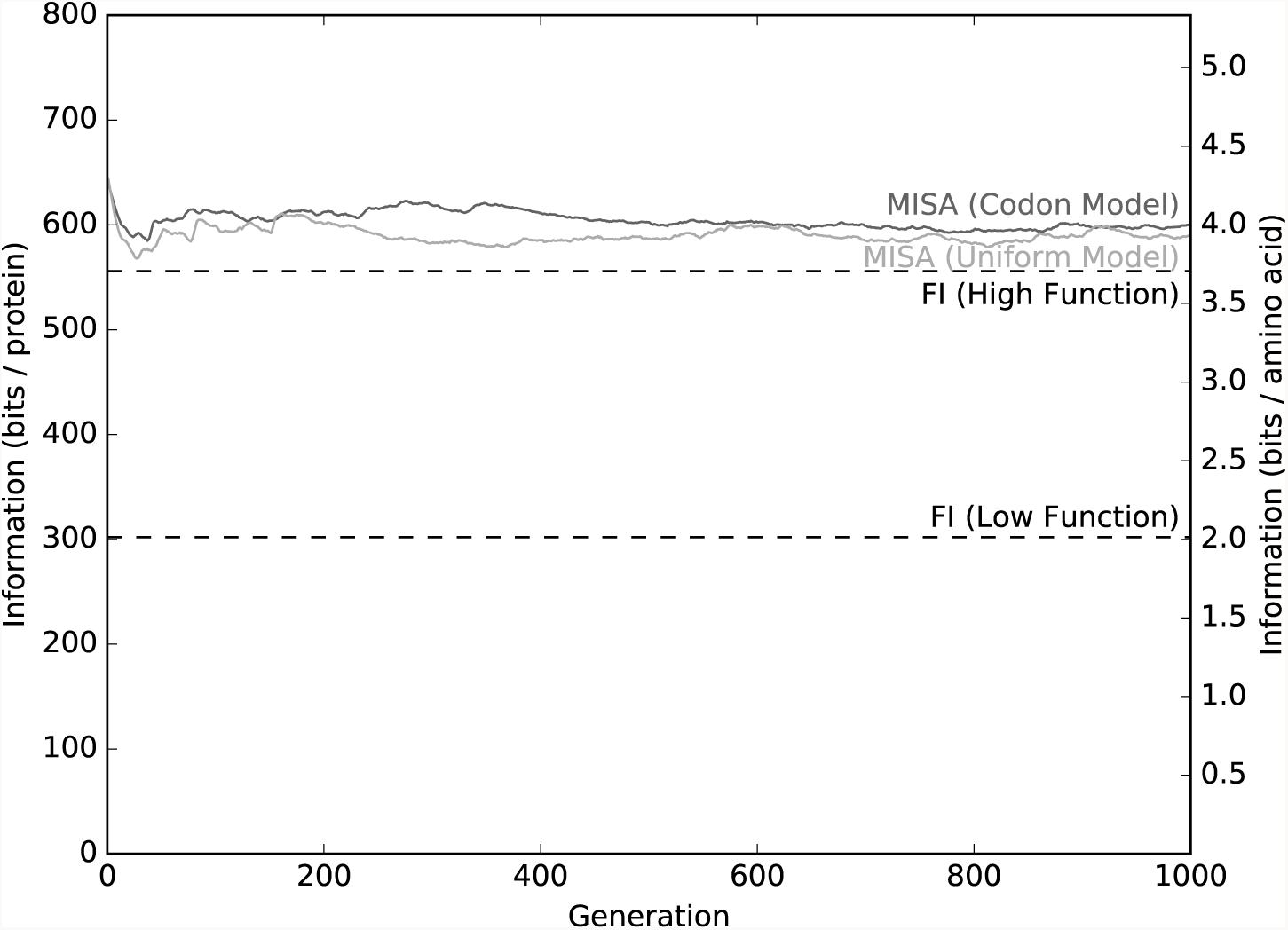
MISA fantastically overestimates the FI of sequences under selection and, therefore, drastically underestimates the number of functional proteins. In this simulation, a functional sequence family is defined as all sequences 150 amino acids long with fewer than 50 amino acids different from a fixed centroid sequence. This space contains 1.8 × 10^104^ sequences (so FI approximately equals 301 bits). A fitness function varies between 50 and 10 differences from the centroid, with sequences closer to the centroid given two times higher fitness than those one mutation father away. All sequences within 10 amino acids of the centroid were defined as highly functional and assigned equal fitness. The FI of the high function group is about 555 bits. The population is initialized to 2,000 copies of a single ancestral sequence exactly 50 amino acids from the centroid sequence, at the edge of functionality. Each generation, ten offspring per sequence are sampled using a mutation rate of 0.01 events per amino acid. The next generation is selected from the offspring with fitness weighted random sampling without replacement. The population converges to a very high level of fitness (Fig. S2), and MISA converges to a value greater than 600 bits, overestimating the true FI by approximately 307 bits, and underestimating the number of functional sequences by about 90 orders of magnitude. We ran this selection simulation using both the codon model of mutations and the uniform model of mutations. Both yield similar results

We have one strong hint that new functions are close to old functions and abundant, and therefore easy to evolve. The same protein family can often perform several different functions. If functionality were astronomically rare and isolated in sequence space, this would be nearly impossible. If the FI argument was even approximately correct, multifunctional proteins and protein families would be extremely rare. Yet we observe them in nature all the time.

Multifunctional proteins are ubiquitous. One excellent example, which our group studies, is the family of P450 enzymes, which metabolize a wide range of different xenobiotic and endogenous molecules in several different ways [1, 20, 27]. Moreover, a large number of individual proteins are multifunctional, with “moonlight” functions that seem unrelated to their first discovered function [14]. The many to many relationship between protein families and functions underscores an important insight. Protein families are defined primarily by our ability to align sequences, but these alignments do not neatly correspond to biochemical function. Individual proteins in most families might be implicated in several functions; many functions can be executed by multiple families. Evolving a specific function may be easy if a large number of proteins and protein families are just a few mutational steps away from the new function.

Quantifying the importance of each evolutionary pathway is a difficult and important scientific endeavor, but beyond the scope of this study. Speaking to the current questions, this data independently tells us the FI argument must be false. This should be no surprise; FI tells us absolutely nothing about the proximity of the new function to extant or ancient families of proteins. So it does not quantify in any way the likelihood of the mutational path to evolve a new function.

Perhaps most surprisingly, in addition to evolving new functions from existing proteins, there is evidence that proteins occasionally arise *de novo* from previously untranslated DNA sequences [2, 24, 26]. In a similarly surprising process, new function can be added to existing proteins by adding protein sequence encoded by previously untranslated DNA sequences too [10]. In all these cases, functional protein sequence in one species corresponds to untranslated DNA in other species. This is a clear sign that functional protein sequences must be common enough to occasionally pop into existence from untranslated DNA.

Once again, FI tells us nothing about how close a new function is to pre-existing and untranslated DNA sequences. While sequence alignments might trace the origin of de *novo* proteins, they prospectively tell us very little about the difficulty of evolving any specific function. The refrain here is the same. Even if we could accurately compute FI, it would tell us nothing about the proximity of new functions to existing sequences. It would tell us nothing about the evolutionary pathways between existing functions and new functions.

## 3 Conclusion

Evolutionary simulations demonstrate that the mutual information of sequence alignments (sometimes referred to as functional sequence complexity) fantastically overestimates the functional information of proteins. This overestimate, in turn, fantastically underestimates the number of proteins with a specified function. Regardless, even if we could estimate functional information, it would tell us very little about the evolvability of new functions. Perhaps most clearly, these simulations demonstrate that FI (as estimated by MISA) is not a “unique property of minds,” but also a signature of shared evolutionary history and non-uniform mutation distributions. For all these reasons, and more, the functional information argument against evolution fails.

## Acknowledgements

The authors thank the MSTP program and the Department of Pathology and Immunology at Washington University in Saint Louis for supporting this work. This work is also suported by the National Library Of Medicine to study Metabolism and Reactivity under award number R01LM012222, and Metabolism in Children under award number R01LM012482.

